# Sexual transmission causes a marked increase in the incidence of Zika in women in Rio de Janeiro, Brazil

**DOI:** 10.1101/055459

**Authors:** Flávio Codeço Coelho, Betina Durovni, Valeria Saraceni, Cristina Lemos, Claudia Torres Codeço, Sabrina Camargo, Luiz Max de Carvalho, Leonardo Bastos, Denise Arduini, Daniel A.M. Villela, Margaret Armstrong

## Abstract

The recent emergence of Zika in Brazil and its association with increased congenital malformation rates has raised concerns over its impact on the birth rates in the country. Using data on the incidence of Zika in 2015-2016 and dengue in 2013 and 2015-16 for the city of Rio de Janeiro (pop: 6.4 million), we document a massive increase of Zika in women compared to men. Even after correcting for the bias due to the systematic testing of pregnant women for Zika, there are 90% more registered cases per 100,000 women in the sexually active age group (15-65 years) than for men but not before 15 or after 65. Assuming that infected men transmit the disease to women in their semen but that the converse is not true, some extra incidence in women is to be expected. An alternate hypothesis would be that women visit doctors more often than men. To test this, we compared the incidence of dengue fever in men and women in 2015 and in 2013 (before Zika reached Rio de Janeiro): in both years, women are 30% more likely to be reported with dengue.

Summing up, women in the sexually active age bracket are far more likely to get Zika than men (+90% increase); sexual transmission is the most probable cause. Women in the 15-65 age group are also 30% more likely to be reported with dengue than men, which is probably due to women being more careful with their health.

## 1. Introduction

Viral diseases transmitted by *Aedes aegypti* mosquitoes, such as dengue, Yellow Fever, Chikungunya and Zika, have been traditionally restrained to the tropical regions of the world, given the vectors intolerance of colder climates^1^. In these regions transmission tends to be modulated by temperature, slowing down significantly when temperatures drop below 20°C. Global warming trends have long been argued to be a threat to public health as it will extend the reach of tropical diseases^1,2,3^. Preparedness for these diseases is the order of the day for the health agencies of countries of temperate climates.

The emergence Zika as a global pandemic threat, is changing the traditional risk scenarios. The Zika virus has the ability to infect other species of mosquitoes^4,5^ thereby potentially extending its reach. *Aedes albopictus*, for instance, is well established in temperate climates. Moreover, the Zika virus can also be transmitted directly from human to human^6^. The sexual route seems to be the most common alternate form of transmission, but the virus is also present in other bodily fluids such as saliva and urine^7,8,9^.

The recent arrival of Zika in Brazil in 2014, and the speed with which it spread throughout the country and into neighboring countries in few months seems to indicate alternate forms of transmission.

In this paper, we look at the age-adjusted incidence of Zika compared to dengue, in the city of Rio de Janeiro, and estimate the relative importance of the sexual route of transmission to the overall incidence of Zika. By considering this additional and asymetric route of transmission, we expect to see a higher incidence of Zika in sexually active women.

## 2. Methods

The data used in this analysis was obtained from the Rio de Janeiro health secretariat, and consists of every case notified of Zika and dengue for the years of 2013 (dengue only) 2015 (Zika and dengue) and 2016 (Zika and dengue, up to april). each record includes date of notification, ICD-10 code, age in years, sex and gestation status.

We also used the officially estimated population of Rio de Janeiro for 2015, based on the 2010 census (6.4 million people, 3 million men, 3.4 million women, figure 1). All age adjusted incidences used in this paper were calculated using this population as standard.

The cases were aggregated to the same age classes as shown in the city’s age pyramid (figure 1). The classes are 5 years wide and incidence values are presented as cases per 100 thousand inhabitants.

Zika incidence in women was calculated with and without pregnant women, to avoid biases. The city’s health services were systematically testing pregnant women displaying skin rash, due to the high risk of babies developing neurological complications caused by intrauterine Zika infection.

To check the statistical significance of the increase in incidence observed in women, we fit a negative binomial GLM model to the number of cases of Zika and dengue(2013) aggregated by age class.

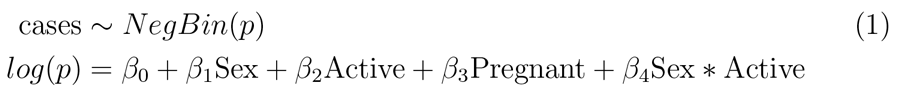

The *Active* dummy variable takes the value 1 for age classes above 15 and less than 65 years of age, and 0 otherwise.

## 3. Results

During the recent Zika epidemic in Rio de Janeiro, which started in late 2015, a total of 29, 301 cases, 20, 315 women and 8, 986 men where notified as suspected Zika cases based on clinical assessment. During the same period 102, 754 total cases of dengue were notified, with 46, 305 being men and 56, 449 being women.

The incidence by age class for Zika in the period of January of 2015 to April of 2016 is shown in figure 2. After removing the pregnant women group from the dataset, we obtain the incidences shown in figure 3. The age distribution of pregnant women removed from the sample is shown in figure 1 of the supplementary material.

For comparison figure 4 shows the age distributed incidence of dengue in the 2015 - 2016 period. Notice that the extra incidence in women is far less pronounced. We also looked at the dengue incidence in 2013 (figure 5), to make sure the pattern is not specific to the 2015 dengue epidemic.

**Figure. 1:**
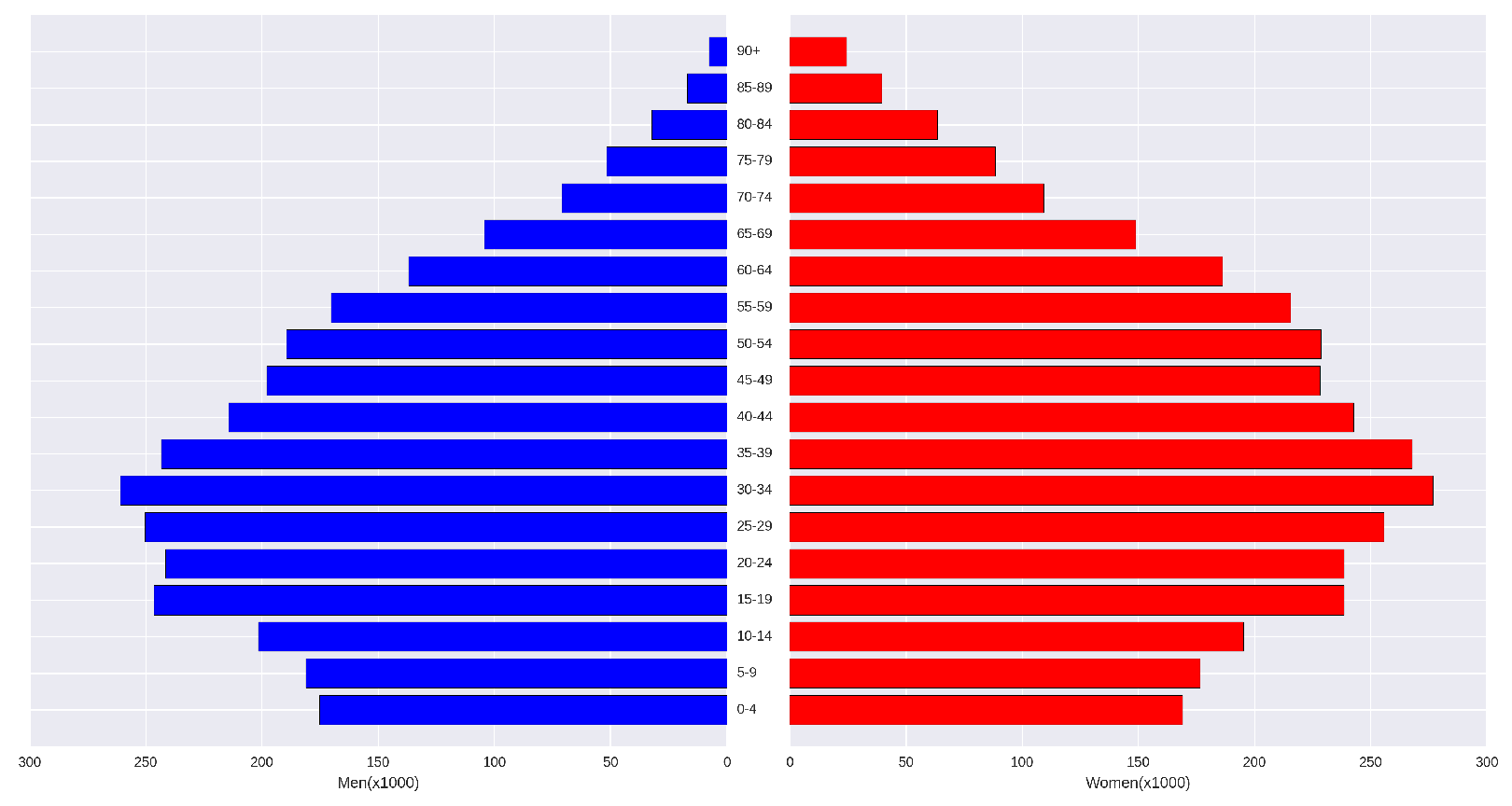
Age Pyramid for the city of Rio de Janeiro. These numbers are official projections based on the census of 2010. The male population is represented in blue, and the female population in red.

The combined incidence in the sexually active age classes, from 15 years old to 65 years old, is shown in table 1 along the ratio of women to men incidence in the same age group.

The regression results point to a significantly higher Zika incidence for sexually active women (1.7767, [0.500, 3.053], *p* = 0.006). Sex alone was not a significant predictor of Zika incidence (−0.2120, [−1.207,0.783], *p* = 0.676). For dengue, being a sexually active woman is not a significant risk factor (0.7196, [−0.321,1.761], *p* = 0.138). Again, sex alone did not prove to be a significant risk factor for dengue.

## 4. Discussion

If Zika is being transmitted both vectorially and sexually in Brazil and other american countries, in the latest epidemic, it is important to estimate the relative importance of each route. According the availabel evidence, the principal way to transmit Zika sexually is through exposure to infected semen. Assuming heterosexual intercourse as far more prevalent than homosexual sex between men, we can expect a surplus of Zika cases in women due to sexual transmission. Moreover active virus have been isolated from semen more than three weeks after the onset of symptoms^9^, which greatly increases the transmission period towards female susceptibles. On the other hand, without significant sexual transmission and in the absence of observational bias favoring women, we would expect equal incidence across age class.

**Table 1.** Aggregated incidence in men and women in the sexually active age group of 15 to 65 years of age. The last column contains the ratio of the women’s incidence to the men’s incidence.

| Disease | Women's incidence | Men's incidence | Ratio(women/men) |
| --- | --- | --- | --- |
| Zika | 5382.58 | 2854.85 | 1.88 |
| dengue 2015-16 | 16628.04 | 13941.05 | 1.19 |
| dengue 2013 | 32201.37 | 25410.52 | 1.27 |

**Figure. 2:**
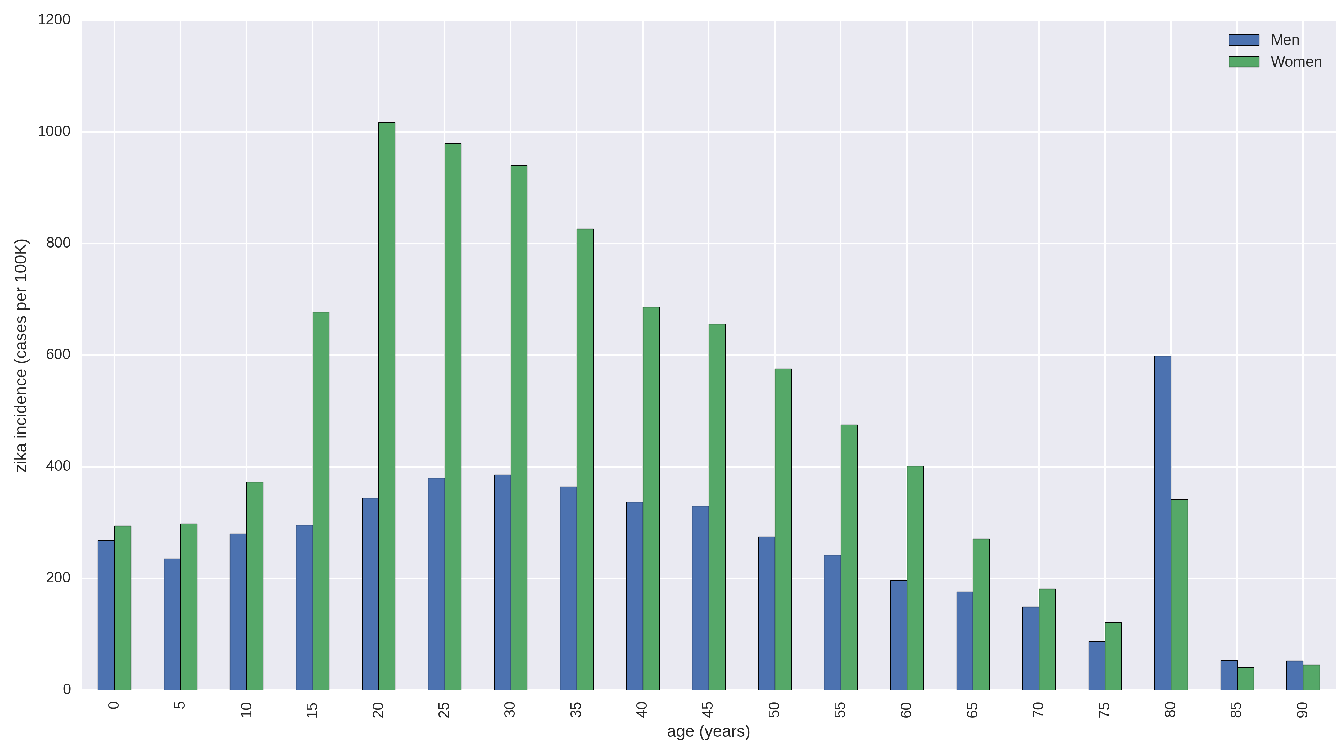
Zika incidence in men and women by age class. Incidence is in units of cases per hundred thousand.

**Figure. 3:**
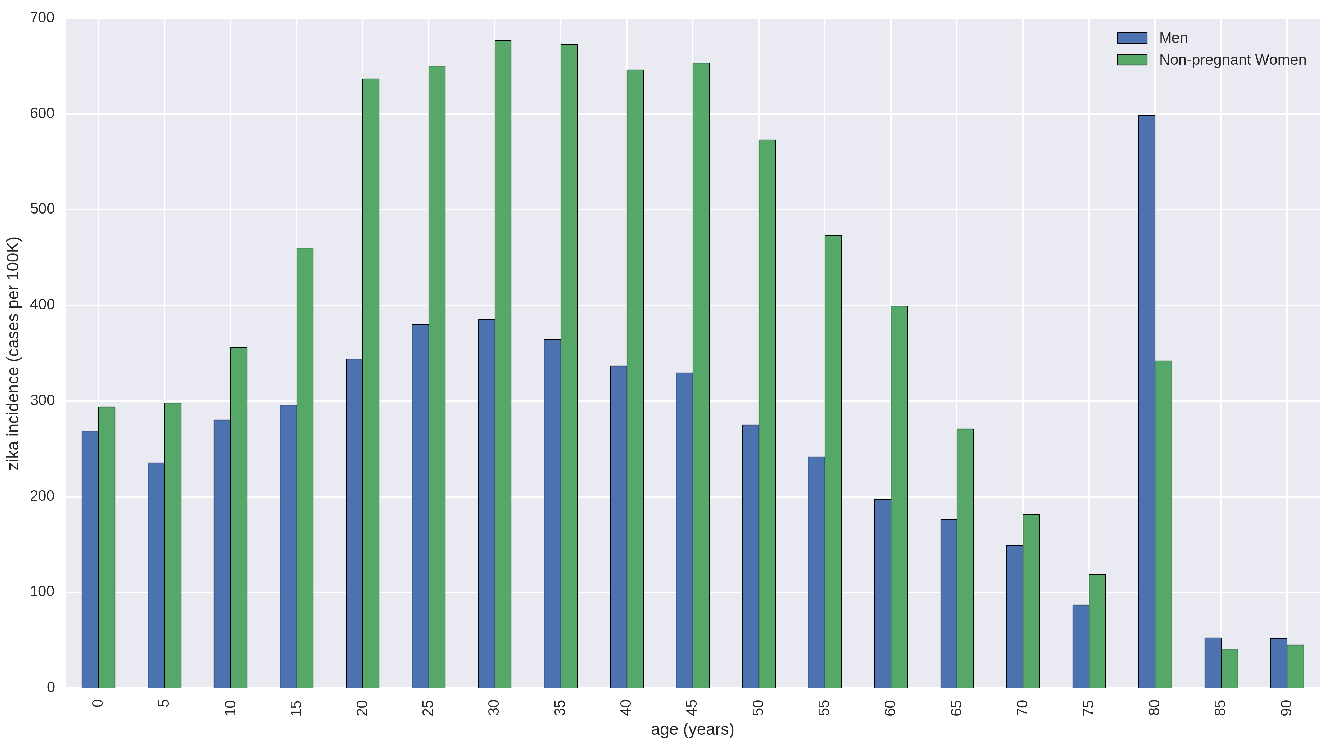
Zika incidence in men and women by age class, excluding pregnant women. Pregnant women are excluded because extra effort was mad by health services to identifiy all possible Zika cases in this group because their babies were at high risk of developing neurological complications.

**Figure. 4:**
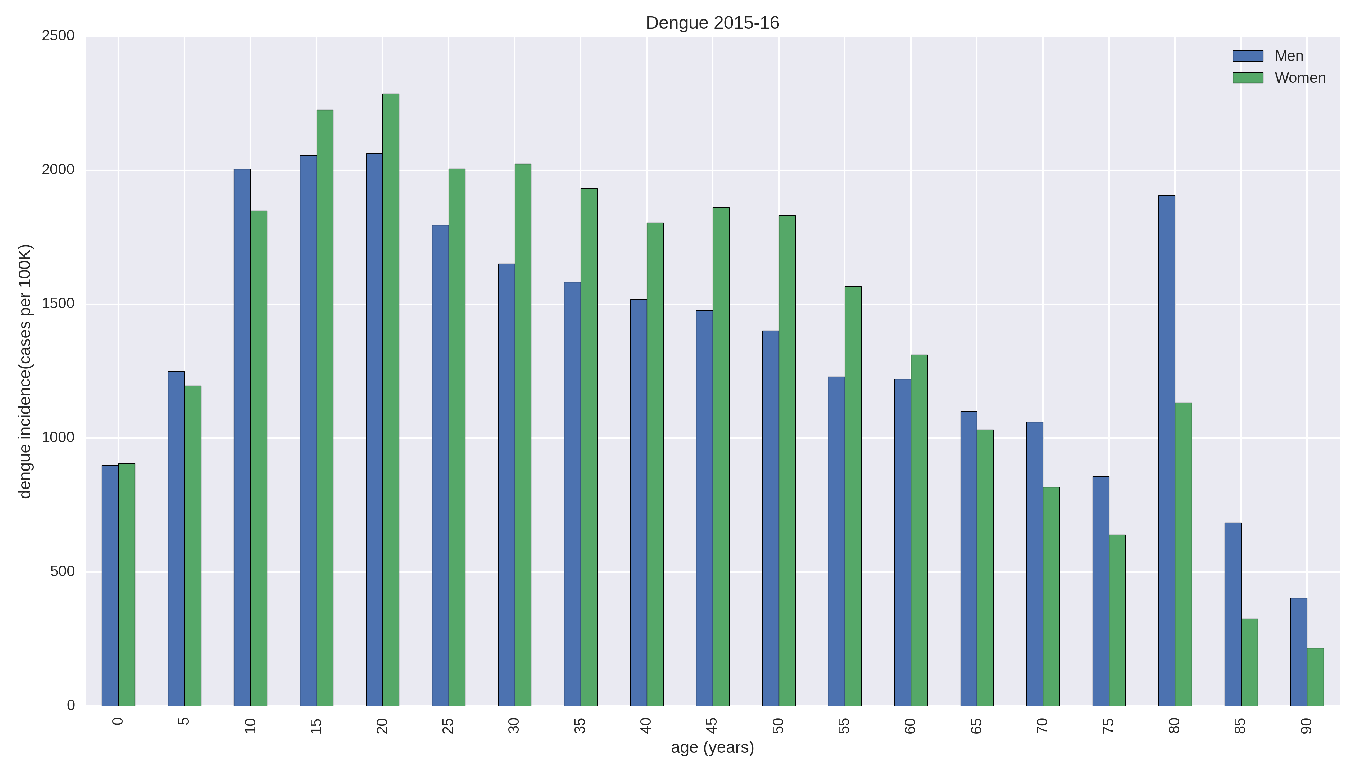
Age-adjusted incidence for dengue in men and women, in the 2015-2016 period.

**Figure. 5:**
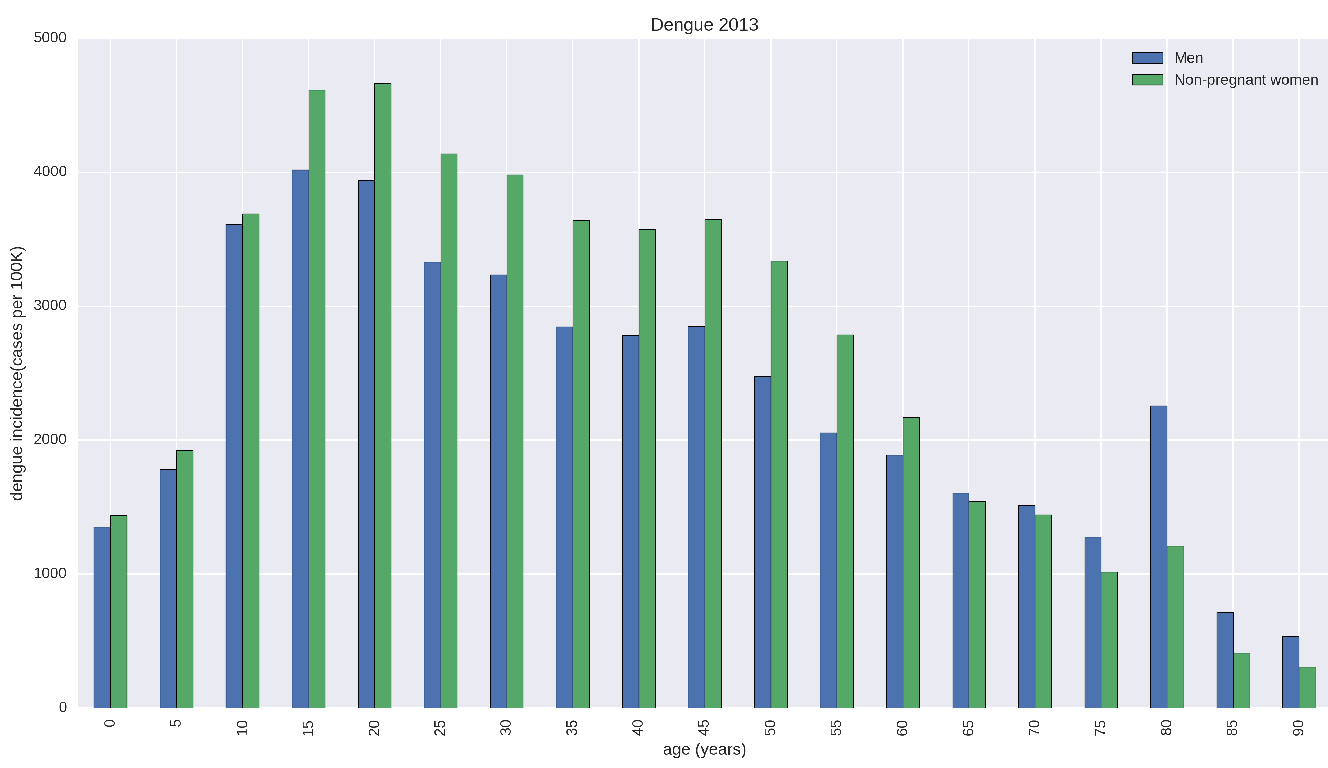
Age-adjusted incidence of dengue in men and women, in the year of 2013. Notice the similar pattern as observed in 2015–16.

What is observed however, is a markedly higher incidence in women as shown in figure 2. As mentioned in the methods section, however, there is at least one observational bias towards reporting women cases, due to concerns about microcephaly and risks to babies. In order to minimize that, we removed pregnant women from the sample (figure 3). What we see is that the extra incidence remains but is quite smaller. But this extra incidence is more pronounced in the reproductive age classes where women are more likely to visit a physician regularly, and thus be detected if they get Zika.

To confirm the hypothesis that women is more likely to be diagnosed with exanthematic fever syndromes during their reproductive years, we looked a dengue incidence by age class (figures 4 and 5). During the 2015-16 period, it is possible that some Zika cases may have been misdiagnosed as dengue, thus “contaminating” the dengue incidence with sexual transmission. So we also looked at the incidence of dengue in 2013, when Zika was unlikely to be circulating in Rio de Janeiro. We also cannot discard that there is evidence that pregnant women are more likely to develop severe dengue^10^. Other factors which could explain the higher incidence in women, are a) women are more likely to stay at home and be more exposed to the vector. b) Women in fertile age, are more predisposed to see a doctor as soon as they develop symptons for fear of complications on an yet undetected pregnancy. We believe that the dengue 2013 data serves a a good control for the differential exposure of women to the vector. As for the behavioral changes associated with the fear of having a microcephaly baby, the data on dengue in 2015-16 also represents a good control for this due to the similarity of the symptoms of dengue and zika.

As can be seen in table 1, the extra detectability of women due to behavior of seeing a doctor regularly, can account for at most 30% higher incidence than men, if we attribute all the extra incidence of dengue in women to this factor. However, for Zika, even discounting pregnant women, we observe an incidence which is almost 90% higher than for men. We attribute this extra incidence to extra cases caused by sexual transmission.

Through sexual transmission, Zika is no longer constrained to tropical and subtropical regions, being able to reach northern Europe, northern United States and Canada, northern Australia as well as Japan and Korea. Although it would be harder for the disease to invade these higher latitudes, the incidences are likely to be higher as men who catch Zika abroad can transmit locally for weeks or even months.

The immediate consequence of this higher incidence of Zika in women in the reproductive age bracket is a much higher expected number of neurologically compromised babies than if the disease was only vectorially transmitted. Going further, women living in Zika infested areas will think twice about falling pregnant, at least those with access to birth control. This could well lead to a drop in the birth rate, particularly in the middle class.

As the Aedes mosquito is known to be present in southern states of the USA around the Gulf of Mexico^3^, in southern Europe^2^ and in northern Australia, an outbreak could start because of a returning traveller, especially if sexual transmission propagates it as well. Health authorities in developed countries are already warning travellers visiting Zika infected areas to consider delaying pregnancies^11,12^. What would happen to the birth rate in these countries if Zika became endemic? A drop in the natality in Europe could have serious economic impacts^13^.

